# Influence of Substrate on Supported Lipid Bilayers: Membrane Pinning, Stretching, Pores, and Remodelling

**DOI:** 10.1101/2025.07.24.666593

**Authors:** Rafael B. Lira, Wouter H. Roos

**Affiliations:** Moleculaire Biofysica, Zernike Instituut, Rijksuniversiteit Groningen, Groningen, The Netherlands

**Keywords:** Model membranes, supported lipid bilayers (SLBs), giant unilamellar vesicles (GUVs), FRAP, FLIM, TIRF, optical tweezers, membrane fusion

## Abstract

Model membranes offer a simplified representation of biomembranes, allowing for controlled structural and functional studies without the inherent complexity of biological systems. Due to the confined space and strong interactions with the support, supported lipid bilayers (SLBs) exhibit markedly distinct properties compared to those of their free-standing counterparts. However, the nano and microscale origins of these differences, structural effects and the related biophysical consequences remain incompletely understood. Using a combination of spectroscopy and micromanipulation techniques, we reveal that SLBs are pinned to the support, creating barriers for lipid diffusion. Furthermore, they are intrinsically stretched and under tension, causing membrane rupture and the formation of stable ∼2 nm aqueous pores that allow the permeation of small hydrophilic molecules. Despite strong adhesion, SLBs can undergo localized shape changes associated with membrane fusion, and membrane nanotubes can be extruded under optical forces. The findings underscore the very distinct properties of model membranes, and we anticipate that they will have important consequences in the field of drug-delivery systems and membrane biophysics.

## INTRODUCTION

Model membranes are invaluable tools in biophysics. They provide simplified, yet versatile and robust systems to study the fundamental properties and behaviour of biological membranes under controlled conditions^1^. In living cells, numerous overlapping processes occur simultaneously, making it difficult to isolate and study a single biological mechanism without interfering with others. In reconstituted systems, specific components (i.e. lipids, proteins, etc.) can be added or removed selectively, and extrinsic variables such buffer medium, temperature, and mechanical forces can be studied separately^2^. Thus, models systems provide a controlled environment to isolate and investigate specific biological phenomena, avoiding the complexity and variability found in living systems.

Among the most popular membrane models are small/large liposomes^3,4^, giant unilamellar vesicles (GUVs)^5,6^, planar supported lipid bilayers^7,8^ and particle/bead-coated bilayers^9^. For simplicity, all supported systems are hereafter simply called *supported lipid bilayers* (SLBs). Liposomes and GUVs possess free-standing membranes bathed in water-based solutions from both sides of the membranes, which provide a symmetric hydration layer that supports large-scale membrane dynamics. In contrast, SLBs are in intimate contact with the underlying support. This brings about important physical consequences. Their leaflets are exposed to highly asymmetric media; the proximal leaflet is in direct contact with the support, whereas the distal leaflet is exposed to the external bathing solution. The space between the proximal leaflet and the support, the so-called *hydration layer*, is a ∼1-2 nm thin space that confers some membrane lubrication^10^. Because it can only accommodate ∼ 3 layers of water molecules^11^, the water molecules in this hydration layer do not behave like bulk water, but instead are under high viscous constrains^12,13^.

Compared to their free-standing counterparts, the presence of an underlying support endows increased mechanical stability to SLBs. On the other hand, it has long been identified that the support also significantly interferes with fundamental membrane biophysical properties, the most well-characterized of which being the increased friction at the proximal leaflet and reduction in molecular diffusion. In supported bilayers, lipid diffusion is 2-10 times slower than in free-standing membranes under otherwise identical conditions^14,15^, although proximal and distal lipids are differentially affected^16,17^. Furthermore, lipids in the proximal leaflet are shown to be pinned at the substrate^18,19^, giving rise to experimentally observed nanoscale hindrances^20^. Other known biophysical consequences of strong membrane-support adhesion include interference with domain formation^21,22^, asymmetric leaflet distribution of charged lipids^23^, and due to the narrow space of the hydration layer, reconstituted proteins exhibit reduced mobility^24^ and are sometimes denaturated^25,26^.

Free-standing bilayers can undergo large shape changes (i.e. deformation, budding, fluctuation) under external stimuli. In contrast, out of plane deformations are highly constrained in SLBs because strong adhesion to the support overcomes bilayer bending energy^9^, thus making SLBs less representative models for membrane dynamics or to study processes associated with membrane remodelling. In principle, weakly-adhered bilayers are amenable to deformation provided they detach from the support. This has been shown in rare cases, such as COPII bud formation upon activation of the endoplasmic reticulum GTPase protein Sar1p^27^, although this represents more of an exception than the rule. Instead, when SLBs are used as a model system for non-planar membranes, they are often prepared on curved patterned substrates, such as buds^28,29^, protrusions^30^, dimples^31^, nanoridges^32^ and nanobars^33,34^. Under external force, fragments of the supported membrane can be locally detached from the support and tubes are extruded^35,36^. This, nevertheless, requires relatively large forces in the order of pN-nN, and while these forces can be overcome by biological remodelling processes such as fusion, it is not clear to what degree the support influences this process. In order to fuse, membranes must overcome several intermediates that are associated with extensive remodelling and that eventually result in the mixing of membrane lipids and aqueous compartments^37^. Given that free-standing and supported membranes exhibit very distinct mechanical properties, we investigated how different their properties are at the nanoscale, and how the ubiquitous biological process of membrane fusion can be recapitulated in the constrained membranes of SLBs.

In this work, we used state of the art spectroscopy and single molecule imaging and manipulation to study the effects of the support on membrane mechanical properties, and probed whether supported membranes are amenable to fusion and remodelling. Spot-variation photobleaching shows that lipid diffusion is reduced in SLBs by a combination of adhesion effects and formation of obstruction barriers. FLIM with environmentally-sensitive molecular probes shows that SLBs are highly stretched, the extent of which depends on the chemical identity of the support, support porosity and membrane phase state. Stretching results in the formation of stable hydrophilic pores of < 2 nm size even in the highly packed membranes in the Lo phase or in the presence of Ca^2+^, two of the conditions known to increase the energy penalty of pore formation^38,39^. Dual-colour FLIM combined with lipid and content mixing assays reveals that charged supported membranes readily undergo both lipid and content mixing. Total internal reflection fluorescence (TIRF) microscopy shows that individual fusion events are extremely fast (under 7 ms), highly efficient (approaching 100%), and complete (full fusion). Notably, the presence of the support enhances fusion efficiency compared to free-standing membranes. Optical tweezers-based force spectroscopy reveals that a high force is associated with fusion and it was used to extrude membrane tethers. The data reveal previously unidentified effects of the support on membrane structure and function that have far-reaching implications, from membrane biophysics, to drug-delivery and advanced sensing technologies.

## RESULTS

### The rationale

The intimate contact, confined space, and limited hydration of the hydration layer alter the structural and dynamic properties of SLBs. The rigid underlying support is expected to constrain membrane remodelling—a fundamental feature of many biological processes, including membrane fusion. Yet, the extent of these restrictions remain unknown. Fusion involves substantial changes in membrane morphology, such as area growth^40^, fluctuation induction^41^, and budding^42^. While fusion has been demonstrated in SLBs^43–47^, most studies focus on molecular determinants like kinetics and regulatory factors, with limited understanding of the geometric aspects of membrane remodelling, including content mixing, area gain, and fusion forces. In this work, we used a diverse range of biophysical tools to gain insights into structural and functional aspects of SLBs and benchmarked them with the properties of their free-standing counterparts.

### Membrane pinning on supported lipid bilayers

The best characterized effect of a support on membrane properties is lipid diffusion. The presence of a solid support has been widely reported to reduce lipid diffusion due to the strong interaction with the proximal leaflet^48–52^. Among the various methods used to study diffusion, fluorescence recovery after photobleaching (FRAP) is a well-established technique. In FRAP, molecules within a defined region are irreversibly photobleached, and fluorescence recovery occurs as unbleached molecules diffuse into the bleached area^53^. The rate of recovery provides a quantitative measure of molecular mobility. In “single-spot” FRAP, diffusion is measured within a defined bleaching area, providing information from the dynamics within that region^54^. In contrast, performing FRAP over areas of varying sizes— known as spot-variation FRAP (sv-FRAP)—allows probing of the structural properties of the environment through which the molecules diffuse^55^. In sv-FRAP, the characteristic recovery time (t_1/2_)—the time at which fluorescence recovers to 50% of its maximum—systematically increases with the size of the bleached area. For freely diffusing molecules, t₁/₂ extrapolates to zero as the bleaching radius approaches zero, while deviations from this behaviour indicate the presence of diffusion barriers or spatial confinement^54,55^.

We performed sv-FRAP on SLBs formed on a glass solid support to measure the diffusion coefficient (D) of membrane lipids and gain insights into potential diffusion barriers. To test for potential charge-mediated effects, we studied neutral bilayers made of DOPC and labelled with the neutral probe NBD-PC (tail-labelled) and negative bilayers made of equimolar amounts of DOPC:DOPG labelled with the anionic probe NBD-PG (tail-labelled). As a reference, we also performed sv-FRAP on GUVs made of the same compositions. Figure 1 A shows typical FRAP recovery curves for increasing bleaching radii on neutral SLBs, demonstrating the slower recovery for larger radii. Figure 1B shows the measured t_1/2_ values for neutral and negative SLBs and GUVs. Note that for all systems, t_1/2_ increases as the bleaching area increases. In GUVs, the presence of charges increases t_1/2_ because of a reduction in diffusion coefficient due to increased viscosity^56^. This effect has been consistently observed in free-standing membranes for several dyes and different experimental methods^40,57^. We calculated D from measurements using the largest bleaching radius, accounting for diffusion during photobleaching (Figure S1). D decreases from 6 ± 1 μm²/s to 3.5 ± 0.6 μm²/s (mean ± s.d.) when the membrane contains 50 mol% charged lipids (Figure 1C). Compared to free-standing membranes, t_1/2_ increases in SLBs for both neutral and charged membranes as a consequence of the reduced D. Interestingly, the measured t_1/2_values for increasing bleaching radii are identical within error for neutral and negative membranes, and demonstrate that in SLBs, friction dominates over charge effects, with measured D ∼ 3 μm^2^/s irrespective of membrane of lipid charge. The reduction in D may be interpreted in two ways: either neutral and negative lipids interact equally strongly with the support, or pinning points create physical barriers that effectively confine lipids into distinct compartments. Because the intercept for SLBs is further away from 0 compared to GUVs, it is likely that the lipids experience some sort of physical constraints, likely due to the presence of confinement generated by pinning points, as sketched in Figure 1D. In summary, the strong interaction of lipids with the underlying support reduces lipid mobility and the presence of pinning points creates physical barriers that additionally constrain lipid lateral diffusion.

**Figure 1.**
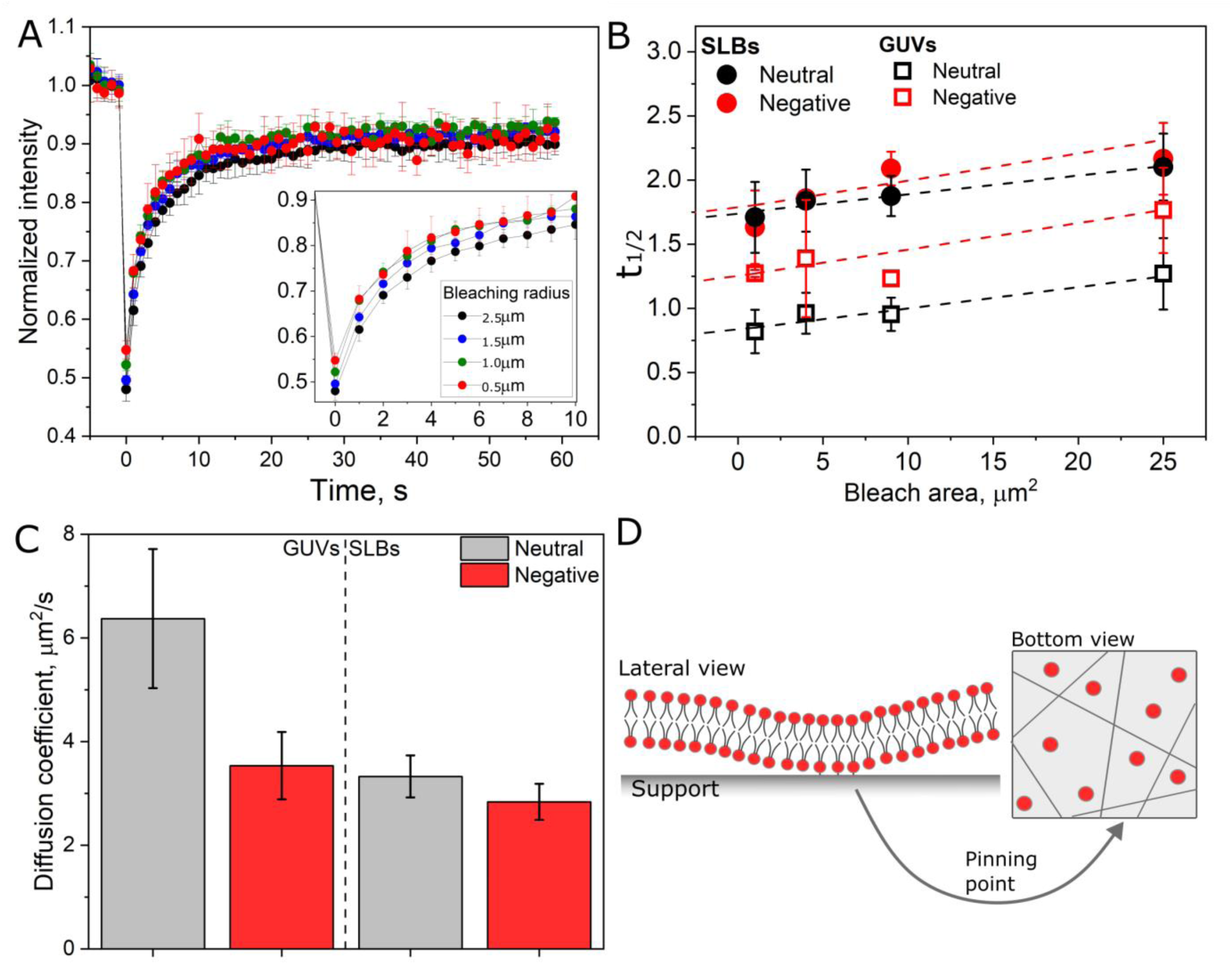
Supported lipid bilayers contain physical barriers that hinder lipid mobility. A, FRAP recovery curves for increasing bleaching areas for neutral (DOPC, labelled with 0.2 mol% NBD-tail) SLBs. The inset is a magnification at earlier time points. Each curve represents the means of 5 independent measurements. B, dependence of t1/2 on the photobleaching area for neutral and negative SLBs and GUVs. Means and s.d. for five independent measurements on each condition are shown. The lines are linear fit and extrapolation to the origin. C, calculated D for neutral and negative GUVs and SLBs measured upon photobleaching with a 2.5 μm radius. D, sketch of pinning in SLBs (left) and how many pinning points create confined areas that hinder lipid mobility, sketched as red circles (right).

### Supported membranes are stretched

We next sought to determine whether the strong interaction with the support influences the membrane’s local microenvironment. To this end, we used the environmentally-sensitive probe FliptR, a molecular flipper probe originally designed to sense membrane tension^58^ and known for its high sensitivity to changes in membrane packing^59^. In loosely packed environments, its fluorescence decays rapidly, predominantly through non-radiative processes, whereas in tightly packed environments, hindered twisting results in slower, radiative decay. We hypothesize that membrane adhesion to a substrate alters membrane packing due to membrane stretching, which can be sensitively detected with FlipTr due to its high dynamic range^60^. In this way, stretching increases the area per lipid and favour FliptR twisting, which is detected by a reduction in lifetime in FLIM experiments. We performed FLIM on GUVs as free-standing bilayers and SLBs formed on planar, solid or porous beads made of polystyrene or silica (Figure 2A). The porous beads have a pore size of ∼ 1-2 nm, which is smaller than the bilayer’s persistence length, and therefore the bilayer formed over is still considered supported.

**Figure 2.**
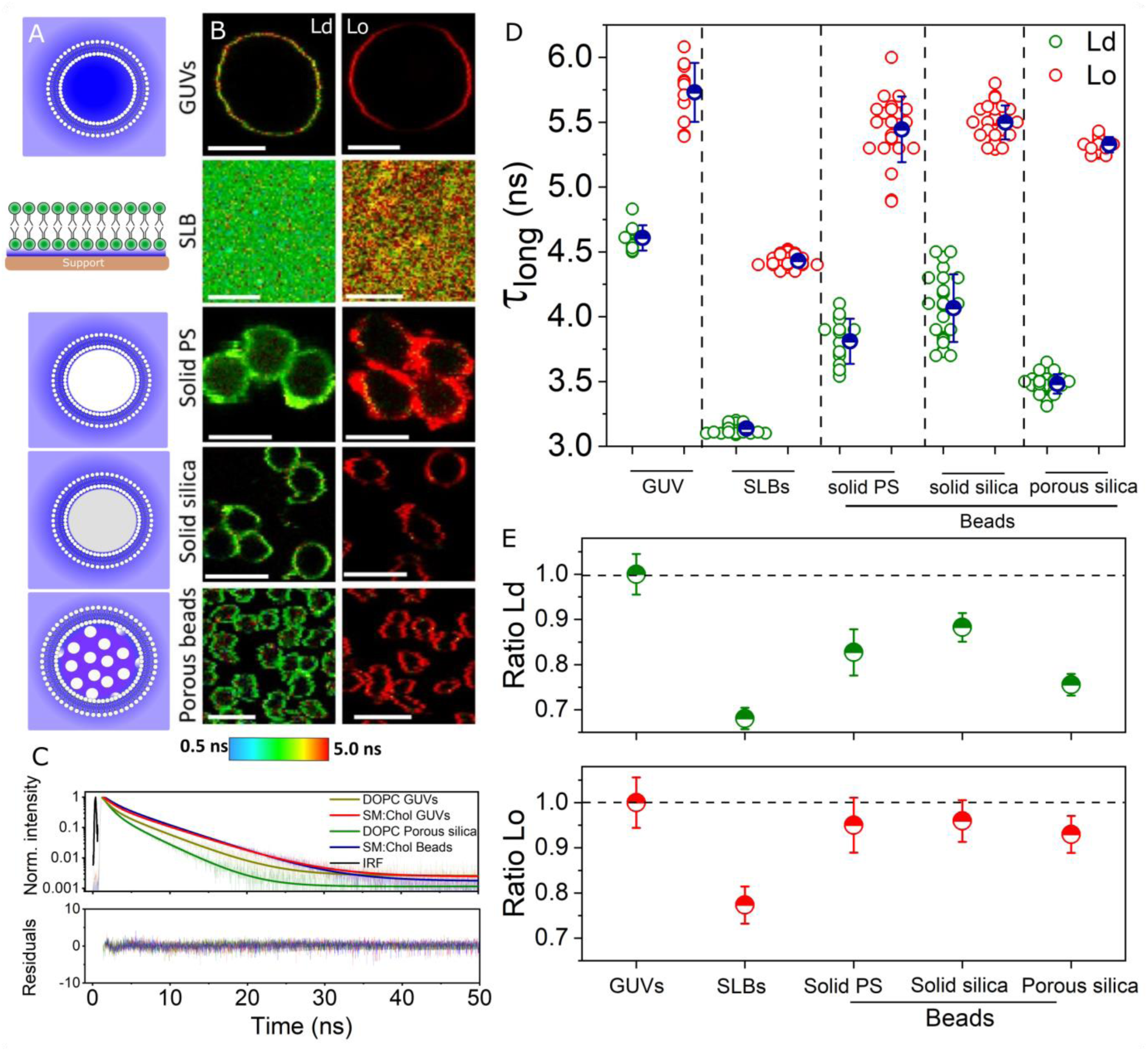
SLBs are stretched. A, sketch of the different membrane systems. The pores are not to scale. B, representative FLIM images of membranes in the Ld and Lo phase labelled with 0.2 mol% FliptR. Scale bars: 5 μm. C, fluorescence decay, fits (two-exponentials) and residuals from the fits for some representative systems. D, measured long lifetime (tlong) for the different membrane systems and phase investigated. Each circle represents a measurement on a vesicle or bead. E, measured lifetime ratio for the various membrane systems in the Ld (green, upper plot) and Lo (red, lower plot) phases. The horizontal hashed line represents the reference values measured on GUVs.

We prepared membranes in the liquid disordered Ld phase (made of DOPC) or in the liquid ordered Lo phase (made of SM:Chol, 7:3 molar ratio), characterized by loosely packed, or highly packed lipids, respectively^61^. Figure 2B shows representative FLIM images of the different systems and membrane phases. For all systems, the membranes are homogeneous at the microscale. Figure 2C shows their respective fluorescence decays, fits and the residuals from the fits. The decays are best fit with a double-exponential, as previously reported^58,60^, in which the long decay (t_long_) is used for environmental sensing. The measured t_long_ for GUVs in the Ld and Lo phase are 4.6±0.1 ns and 5.7±0.2 ns, respectively, in agreement with prior reports^58,60^. Strikingly, t_long_ is significantly reduced for all supported bilayers in the Ld phase, and the magnitude of reduction depended on the support type. The greatest effect is observed for SLBs formed on glass (t_long_ 3.1±0.1 ns). Due to their high cohesion and stretching elasticity^62^, Lo-phase SLBs are expected to be less sensitive to changes, which was indeed the case. However, even for these membranes, the effect observed in Lo-phase SLBs formed on glass is so pronounced that their fluorescence lifetime (t_long_ 4.4±0.1 ns) is comparable to that of Ld-phase free-standing membranes. These effects become clearer when we plotted the ratio of lifetime for a given membrane system relative to the reference values measured GUVs (Figure 2E). For all supported Ld membranes, lifetime ratio is significantly lower, whereas those measured in supported membranes in the Lo are less pronounced, except for supported bilayers formed on planar support, which shows the strongest effects.

Because a decrease in lifetime is an indication of free probe twisting, we attribute these effects to membrane stretching^58^. Interestingly, the lifetime measured on SLBs formed on solid silica more closely resembles that observed on solid polystyrene beads, rather than on porous silica beads, highlighting the influence of support porosity. Notably, nanoscale roughness has been linked to reduced membrane mobility, consistent with our observations^63^. In summary, our findings reveal that strong interactions with the support not only lead to the well-documented reduction in lipid diffusion, but also induce significant stretching of the adhered bilayer, causing structural alterations even in the highly ordered and stable Lo phase membranes.

### Supported membranes are permeable to small molecules

Highly stretched membranes are more prone to rupture^64^. To assess whether the increase in tension has an impact on membrane integrity, we bathed GUVs and membrane-coated solid polystyrene or porous silica beads in a solution containing the small water-soluble fluorescent probe KU530 (< 1 nm hydrodynamic diameter), a green-emitting molecule that exhibits a long fluorescent lifetime^65^. The pore size in the porous beads is ∼ 2 nm, much smaller than the bilayers persistence length^66^, and therefore the bilayer over the pore is considered supported, in contrast to pore-spanning bilyaers^67^. In intact membranes, KU530 should be retained in the outer medium, whereas if the membrane contains pores, the probe will be able to permeate and enter the GUVs or beads provided the pores are larger than probe’s hydrodynamic radius. The membranes are labelled with the green probe FlipTr, and membrane and leakage probes are resolved based on lifetime differences despite similar emission using FLIM. As shown in Figure 3A, GUVs made of DOPC are not permeable to KU530, in agreement with several other reports for many other dyes of comparable sizes^68–70^. If we encapsulate a fluorescent dye in the space between a solid bead and the membrane, we do not observe a detectable signal (Figure 3B) because this space is too small to accommodate a sufficiently high probe concentration. Hence, under our experimental conditions, solid beads are not suitable for membrane permeability studies. When KU530 is added to SLBs formed on porous silica beads, the probe signal is observed in the beads (Figure 3C). For the bead shown in C, FlipTr lifetime measured on the membrane is 4.6±0.1 ns, identical to reported values on DOPC membranes^58,60^, whereas the KU530 lifetime measured in the bead interior was 18.0±0.1 ns. Note that KU530 has an intrinsic affinity to the silica material, resulting in its accumulation (and hence stronger signal) in the bead interior, an effect observed for other probes and at different pHs (Figure S2). The results show that bilayers formed on porous beads are permeable to small dyes.

**Figure 3.**
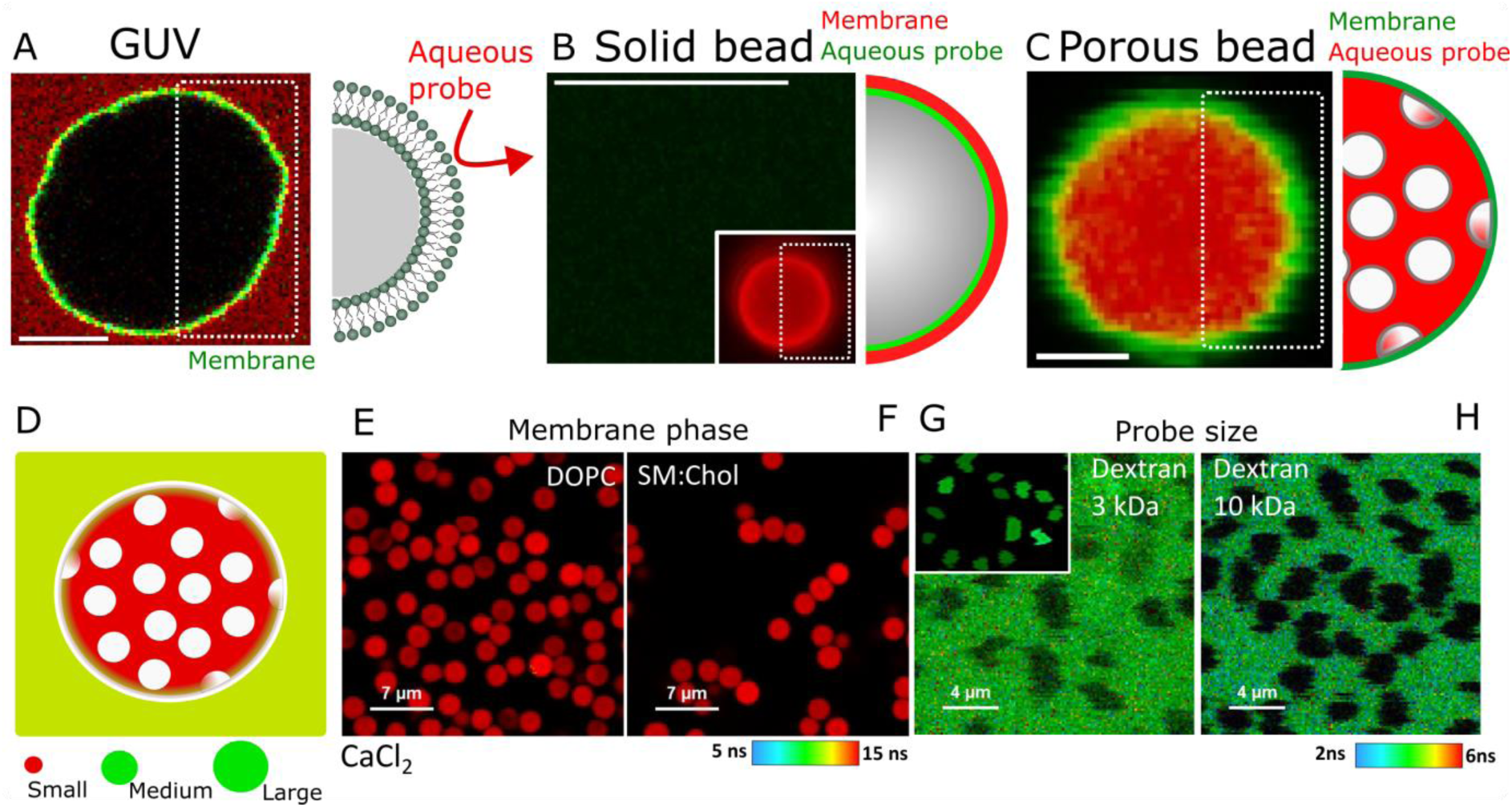
Supported lipids bilayers are permeable to small molecules. A, a representative DOPC GUV incubated in the presence of 0.02 mg/ml KU530 (red). The membrane is labelled with FliptR (yellow). Both probes are green-emitting and they are resolved in the time domain using FLIM. On the right, a sketch of probes being unable to permeate through free-standing membranes. B, DOPC membranes formed on solid polystyrene bead. Rhodamine 6G (5 μM) encapsulated in the aqueous space (green signal in the sketch on the right) is not detectable. The membrane is formed as observed by the red signal in the inset. C, membranes formed on porous silica beads are permeable to KU530 (green signal inside the beads). The right sketch represents the dye inside the beads (not to scale). Scale bars A-C: 5 μm. D, sketch of porous silica beads SLBs incubated with probes of different sizes. E and F, SLBs in the Ld phase (DOPC) and Lo phase (SM:Chol, 7:3 mol ratio), respectively, formed on porous silica beads in the presence of 5 mM CaCl2. G and H, SLBs formed on porous silica beads incubated in the presence of 0.1 mg/ml Dextran 3 kDa and 10 kDa, respectively. The inset in G shows the sample after > 1h incubation. The lifetime colour code is shown for the different probes. The beads in Figure E and F are better defined (not blurred) due to membrane adhesion to the glass slide in the presence of CaCl2. Scale bars in A and B: 7 μm. Scale bar in C: 1 μm.

Stability of aqueous pores in membranes depends on edge tension (γ), with higher γ values favouring pore closure^71^. We have recently shown that membrane γin PC GUVs is high, in the order of 40 pN^38^, resulting in fast closure of formed pores within ∼ 50 ms^68^. Edge tension is further increased in the presence of Ca^2+^ ions^38^, as well as for highly viscous membranes in the Lo phase^72^, with γ ∼ 200 pN and an almost instant pore closure (within ∼ 10-20 ms)^39,72^. Even in these conditions, the bilayers formed on porous beads are permeable to KU530 (Figure 3D-F). Lastly, we probed the size of the formed pores by studying permeation to water-soluble fluorescent probes of increasing size. When we added the medium-sized probe Dextran 3 kDa (∼ 1.5 nm hydrodynamic diameter^73^), some degree of permeation is already observed upon mixing, and accumulation in the beads is observed at longer incubation times (> 1h). The larger probe Dextran 10 kDa (∼ 1.9 nm^73^) was retained outside even at prolonged incubation (> 1h) – Figure 3H. We conclude that the pores formed in supported membranes are small (< 2 nm) and stable despite the high edge tension of highly packed membranes and in the presence of Ca^2+^. Hence, the high edge tension that confers free-standing membranes self-healing capacity is not sufficient to repair defects in stretched supported membranes.

### Free-standing and supported membranes are amenable to fusion

SLBs are typically strongly adhered to their underlying support, with adhesion strengths ranging from 0.1 to 1 mN/m^74^, potentially inhibiting membrane remodelling processes. We next sought to determine whether, despite strong adhesion, SLBs are also amenable to fusion in a similar manner to that observed in free-standing bilayers. Membrane fusion has been demonstrated between membrane-coated beads^75,76^, between coated-beads and planar support^45,77,78^, as well as between vesicles/enveloped viruses and planar membranes^43,46,47^. This indicates that, despite adhesion, membranes retain the ability to bend at the microscale and/or nanoscale. However, due to the solid nature of the support, the distinction between lipid and content mixing is cumbersome and, to the best of our knowledge, content mixing has not been demonstrated with membranes formed on a solid (planar or bead) support.

To study membrane fusion, we developed a two-channel FLIM assay to simultaneously probe lipid and content mixing. In this assay, cationic liposomes labelled with the far-red membrane probe DOPE-Atto647 and encapsulating the green aqueous probe Dextran 10 kDa were incubated with target membranes labelled with the green dye FliptR. Lipid and content mixing are observed in two spectral channels, whereas the green dyes Dextran 10 kDa and FlipTr are resolved based on their lifetime difference. Lipid mixing without content mixing is a hallmark of hemifusion^79^. Figure 4A shows the experimental setup. After incubation with liposomes, lipid mixing is detected by the appearance of DOPE-Atto647 signal in the target membrane, and content mixing is detected as Dextran 10 kDa signal in the lumen of the target membrane. As shown in Figure 4B (i), both lipid and content mixing are detected in free-standing GUVs, showing full-fusion between the liposomes and GUVs, in agreement with previous observations^72,80^. When the liposomes were incubated with membranes formed on solid beads, a weak yet clear lipid mixing signal is observed. This confirms that SLBs can undergo fusion. However, as expected for solid beads, we are unable to resolve whether full fusion takes place as the content signal cannot be spatially resolved in the hydration layer (Figure, 4B ii). Strikingly, liposome incubation with SLBs formed on porous beads results in lipid mixing accompanied by a clear transfer and retention of Dextran 10 kDa in the bead interior (Figure, 4B iii). Figure S3 shows that the fluorescence signals can be clearly resolved based on lifetime differences for all systems. Simultaneous lipid and content mixing of liposomes with SLB formed on porous beads are also observed with other dye combinations (Figure S4).

**Figure 4.**
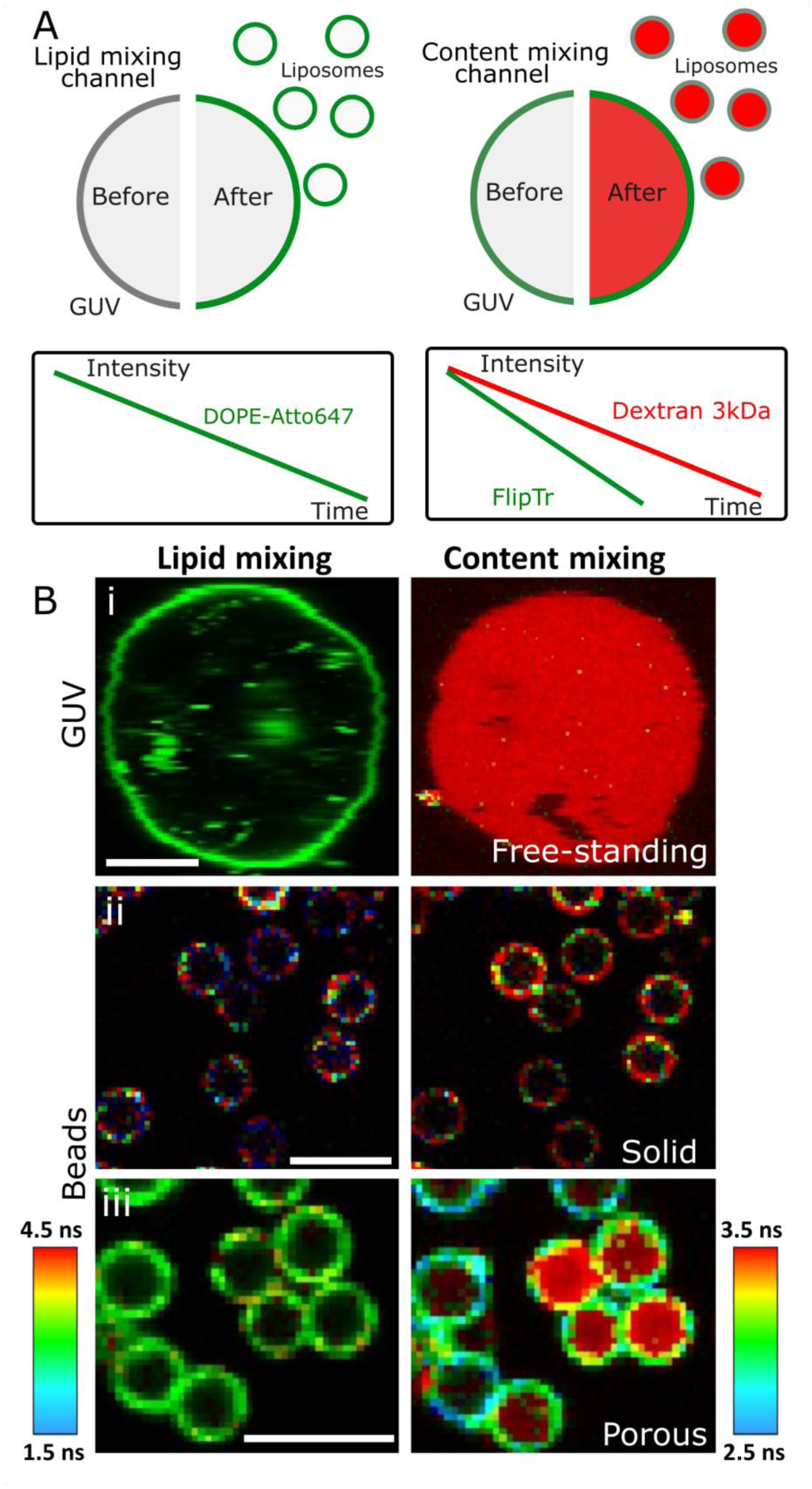
Simultaneous lipid and content mixing in model membranes. A, sketch of the two-colour FLIM experimental setup. In the lipid mixing channel, non-labelled acceptor membranes (GUVs or beads) attain a fluorescence signal at the membrane upon lipid mixing with fusogenic liposomes labelled at the membrane (green). In the separate content mixing channel, fusion of liposomes encapsulating the aqueous probe Dextran 3 kDa (red) with labelled (FliptR, green) or unlabelled acceptor membranes results in content mixing. Membrane and content signal are temporally resolved due to their distinct fluorescence lifetimes (sketch). B, representative images of free-standing GUVs (i), solid (ii) or porous (iii) beads incubated with the fusogenic liposomes. The colour scale shows the lifetime range in both channels. Scale bars: 5 μm.

In addition to fusion between (free-standing) liposomes and SLBs formed on solid or porous beads, membrane fusion can be observed in all other configurations (Figure S5 and S6). Figures S5A-S5C show the fusion between neutral and cationic membrane-coated beads. Fusion is confirmed by bead binding and lipid mixing. Some of the binding pairs do not undergo lipid mixing (Figure S5 B), in agreement with the relatively weak interaction and inefficient fusion observed between neutral and cationic membranes^40,72^. If both membranes are electrically neutral, they do not interact nor fuse (Figure S5 D-E). Conversely, fusion as observed via content mixing, also occurs with supported bilayers formed between cationic and anionic porous beads (Figure S6). In summary, supported membranes, like their free-standing counterparts, support lipid and content mixing, demonstrating their capacity to undergo the microscale remodelling processes necessary for full fusion. However, only vesicles or porous beads are amenable to both lipid and content mixing.

### Supported bilayers enhance fusion and suppress large-scale membrane deformations

To uncover the molecular mechanisms and kinetics of membrane fusion, we examined the process at the level of single events using total internal reflection fluorescence (TIRF) microscopy to visualize individual liposomes fusing with a planar lipid bilayer in real-time. TIRF generates an evanescent excitation that is confined to a few hundred nanometers in the axial direction. Thereby background fluorescence is reduced and the spatial resolution is enhanced while maintaining the high temporal resolution typical of wide-field imaging^81,82^. This enables detection of fusion intermediates and kinetics with high-throughput (Figure 5A). Figure 5 B represents the expected intensity changes for each of the fusion events when observed with lipid mixing only. Incoming fusing vesicles flatten and approach the surface, resulting in a distinctive fluorescence burst that marks the fusion event^46,81,83^. At the single-vesicle level, docking is observed as the appearance of a diffraction-limited spot. Hemifusion is characterized by a burst that decays to approximately half of its intensity. Full fusion is marked by a fluorescence burst that decays to background. Conversely, transitions (from docking to fusion, or from hemifusion to full fusion) can also be resolved. If the liposomes also entrap an aqueous fluorescent probe, lipid and content mixing can be observed simultaneously.

**Figure 5.**
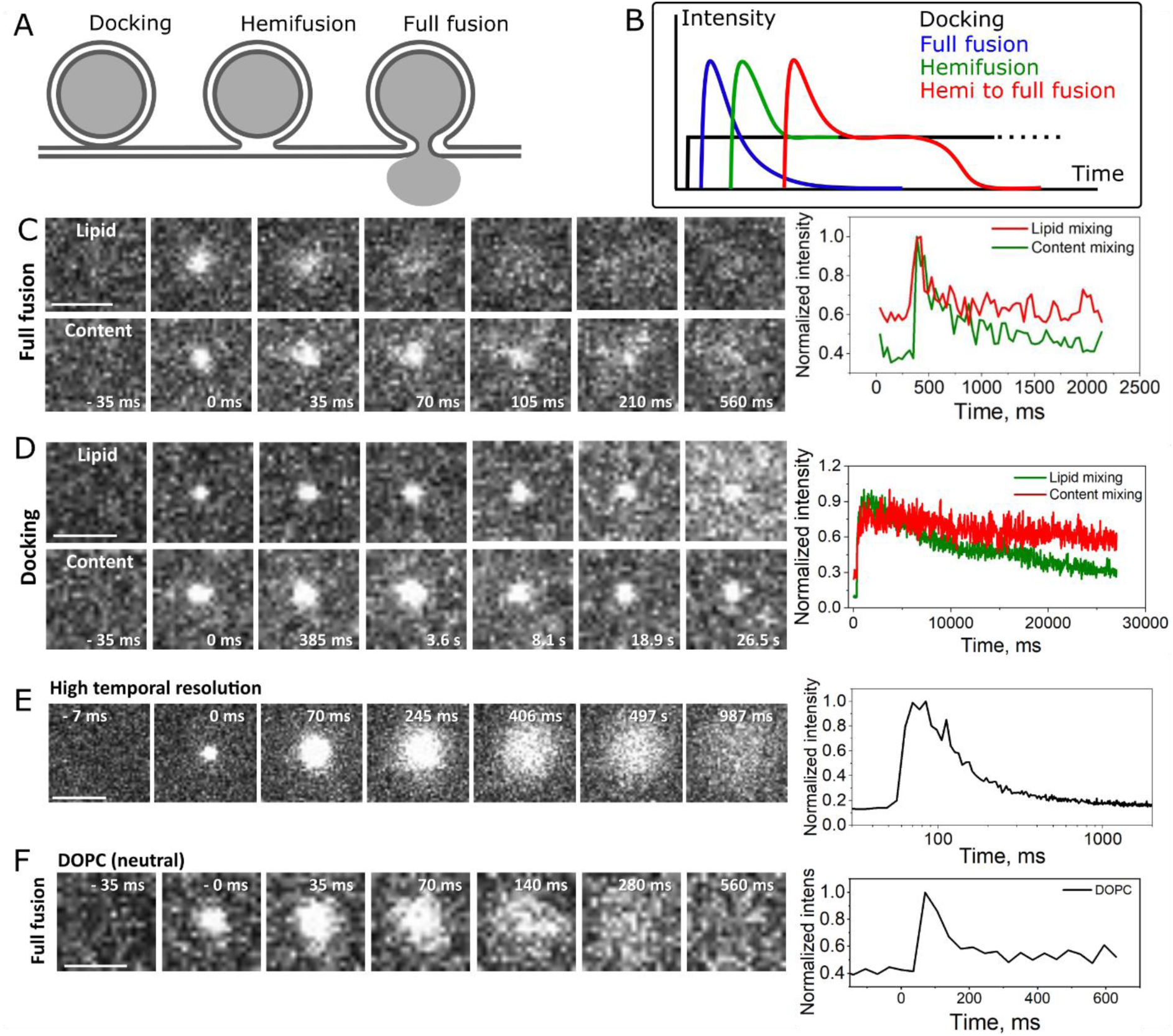
Single fusion events are extremely fast and complete. A, sketch of the assay. Incoming liposomes may undergo different types of interactions with the planar SLB, such as docking (no mixing of lipids), hemifusion (mixing of outer leaflets) and full fusion (mixing of outer and inner leaflets). If an aqueous probe is encapsulated in the liposomes, full fusion results in probe release under the membrane. B, expected changes in fluorescence intensity for the different types of fusion events. C and D, two-colour TIRF to probe lipid and content mixing for a liposome undergoing full fusion and docking, respectively. The intensity traces are shown in the plot on the right. Note the distinct timescales. E, a full fusion event (lipid mixing) observed under 7 ms/frame. F, a neutral liposome made of DOPC undergoing full fusion with the planar SLB. Scale bars: 2 mm. The times refer to the onset of fusion.

Figure 5C shows representative snapshots of a single liposome labelled with DOPE-Atto647 and encapsulating the aqueous probe sulforhodamine B and undergoing full-fusion with the planar SLB. The intensity time traces in both channels are also shown. The appearance of a diffraction limited spot is very quickly followed by fusion, resulting in a diffuse cloud of probes as (i) the lipid probe diffuses in the membrane (lipid mixing) and (ii) the content marker diffuses into the hydration layer. Fusion is extremely fast and occurs within one frame (at 35 ms/frame). In contrast, stable docking results in the appearance of a diffraction-limited spot in both channels that remains stable during the course of the experiment (except for photobleaching observed in the lipid mixing channel), Figure 4D. Considering that liposome fusion releases the entrapped cargo molecule in a quasi-two dimensional space of 1 nm height (h)^10^ and infinite width, the time (t) it takes for the probe to laterally diffuse is t = /4D, where D is the probe diffusion 400 μm^2^/s^84^ is ≈ 0.6 ns. This is much faster than the observed half-time fluorescence decay of ∼ 5 ms, and indicates that the diffusion of aqueous probes is hindered within the hydration layer.

High-speed TIRF imaging at 7 ms per frame reveals that fusion events occur within a single frame upon vesicle docking (Figure 5E). This indicates a fusion rate as fast as, or even faster than, any previously observed single fusion event in vivo^85,86^ or in vitro^87,88^. Notably, >95% of all observed events (i.e. fusion efficiency) correspond to full fusion (Video 1). Fusion occurs only in the presence of a net charge: an excessively high concentration of liposomes fusing with the SLB depletes its charges, effectively halting further fusion (Video 2). Additionally, while fusion in free-standing GUVs induces membrane budding due to curvature effects^40^, no such deformations are observed in SLBs. This suggests that although the membrane remodelling required for fusion can take place in supported bilayers, the strong adhesion to the support suppresses large-scale membrane deformations.

In free-standing GUVs, full fusion requires high charge density^56^, whereas fusion with neutral GUVs is slow, inefficient and primarily occurs through hemifusion^40^. Figure 5F shows that neutral liposomes undergo full fusion with cationic SLBs within one frame (35 ms/frame). Similarly to fusion of negative liposomes, nearly all neutral liposomes fuse (i.e. ∼ 100 fusion efficiency) within a few milliseconds (Video 3). Therefore, under conditions where liposome fusion with free-standing membranes is inefficient, supported lipid bilayers promote highly efficient full fusion with unprecedentedly fast kinetics. This demonstrates that stretched SLBs favour fusion, consistent with the stimulatory effects of membrane tension on fusion observed in other systems^64,89^. In summary, strong interactions with the support and increased membrane tension promote fusion; however, these same interactions suppress the large-scale deformations typically observed in GUVs.

### Membrane fusion interaction forces in supported membranes

One of the main advantages of studying membrane fusion with supported bilayers is to controllably manipulate membrane-coated beads under optical forces using optical tweezers. Together with AFM^77,90^, this is one of the few methods able to measure contact forces of fusing membranes^91^. We used a method that combines microfluidics, force spectroscopy and fluorescence imaging to study the interaction forces between bilayer-coated beads upon membrane fusion. Bilayer-coated polystyrene beads were manipulated with subnanometer precision and brought in contact by moving one bead towards a second, stationary bead where the trapping force was measured with a 0.1 pN precision^92^. Previously, this has been used to study the interactions and fusion of coated beads in the presence of calcium-sensitive proteins Doc2b^76^ and the C2AB fragment of synaptotagmin^93^. In the presence of calcium, these proteins mediate interaction forces in the range of several hundreds of pN, and the force was interpreted as the formation of a fusion stalk (i.e. hemifusion)^76,93^. Upon bead retraction, the appearance of a lipid tube connecting the membranes could be observed. Apparently, single tubes were occasionally formed and their forces were detected until the force dropped to zero, indicating the loss of tube connectivity^91^.

To gain insights into the forces associated with charge-mediated membrane fusion, we performed force spectroscopy using optical tweezers. A microfluidic chamber was used to controllably capture specific interacting pairs, and the interactions and fusion were imaged using confocal microscopy. As schematically shown in Figure 6A, cationic beads labelled with DOPE-Rh (green) and anionic beads labelled with DOPE-Atto647 (red) were loaded on separate microfluidics channels. Upon trapping one of these beads in one channel, we quickly moved to the next channel to capture a beads of opposite charge while the SLB composition is identified from confocal microscopy (Figure 6B). We bring the coated beads into contact by moving the right-hand bead toward the stationary left bead, measuring the detection force exerted on the left bead, and recording force-distance (F-D) curves to derive the interaction forces. Typical contact forces are 10-30 pN and the beads are left to interact for 30-60 s (Figure 6 C). For non-interacting beads, the measured force goes back to 0 upon retraction. For the bead pair shown in Figure 6D, a linear increase in force to a maximum of 100 pN is detected upon retraction with a negligible increase in bead distance, until an abrupt force drop, where the beads separated. The force does not drop to zero, but instead to an intermediate value. As retraction progresses, additional force drops are observed (Figure 5D, inset). The fact that the force is significantly above zero at large separation distances indicates that the membranes are connected (i.e. via membrane nanotubes), as previously reported^76,93^, as indeed observed from confocal imaging. In some approach-retraction experiments, multiple tubes are observed (Figure S7). Importantly, the generated tube as well as the beads are labelled with both dyes, a hallmark of lipids mixing. With further separation, the force drops to zero as the connectivity is lost. Upon a new approach-retraction attempt, no further interaction is observed, likely due to charge neutralization. Of the 22 pairs that exhibited interaction forces, 5 could not be separated —even at forces approaching the ∼500 pN limit of our optical trap—causing the beads to escape from the trap during retraction attempts. These strong, irreversible interactions are consistent with previous observations^76,93^. These interactions are charge-specific; approach-retraction attempts of beads of similar charges do no result in detectable forces (Figure S8). The results highlight the strong interaction forces driven by electrostatic interactions and the formation of multiple membrane tethers during mechanical retraction, and that lipid tubes could be extruded in fused membranes formed on supported membranes.

**Figure 6.**
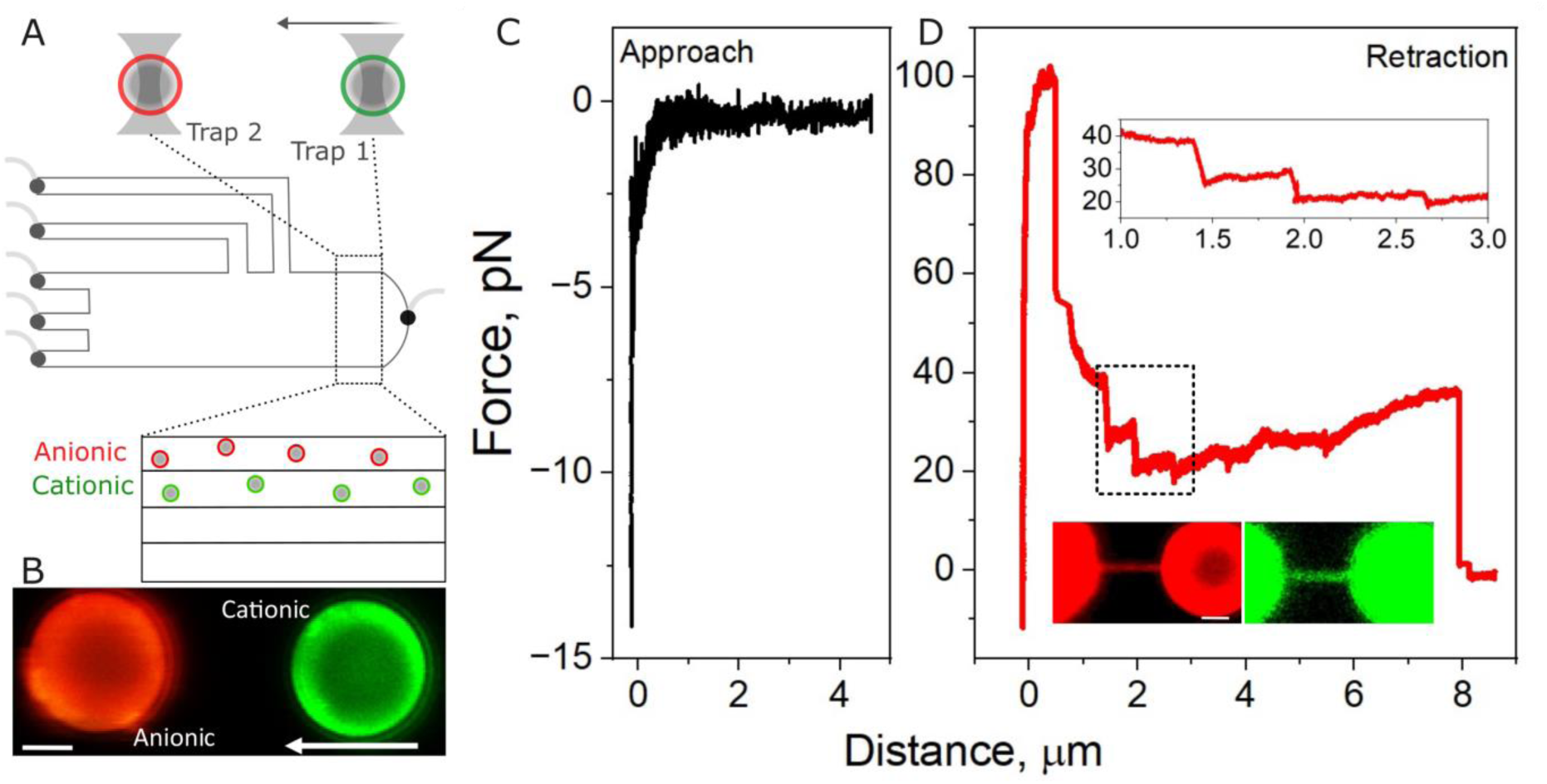
Forces associated with charge-mediated membrane fusion. A, sketch of the beads trapped in a microfluidic device. A double-trap is used to controllably move the bead on trap 1 towards the bead on trap 2, where the force measurements are recorded. Anionic (red) and cationic beads (green) are loaded in different microfluidic channels. B, representative confocal snapshot of two optically trapped beads. The composition of their membranes is identified based on their colour. Scale bar: 2 μm. C, measured force in the approaching phase. D, measured force upon bead retraction. The inset graph is the zoom-in of the region marked in the hashed square. The images below show the separate red/green channels, demonstrating the presence of a lipid tube as well as lipid mixing.

## DISCUSSION

Membrane models are invaluable tools in biophysics, offering a close mimic of the composition, structure and biophysical properties of biomembranes. Among which, SLBs offer a stable geometry, are ideal systems for characterization with surface-sensitive techniques, while endowing the membrane enhanced mechanical stability. However, SLBs are also under the physical constraints of their support, which has shown to affect molecular diffusion, friction and overall remodelling capabilities, all of the features that are unconstrained in their free-standing counterparts. In this work, we show that, in addition to these effects, SLBs are also pinned to the surface, creating physical barriers for lipid diffusion, they are stretched and under tension, resulting in the formation of small (< 2 nm) and stably open pores. The magnitude of stretching depends on the nature and porosity of the underlying support, and in extreme cases (i.e. in bilayers formed on planar glass slides), the highly packed membranes of the Lo phase are stretched to a point where lipid packing is compared to that of fluid membranes in the Ld phase. Elevated membrane tension facilitates fusion even under conditions where it is otherwise inefficient in free-standing membranes, with high associated fusion forces. However, the strong adhesion to the support restricts the large-scale membrane deformations observed in free-standing systems, although membrane fragments (i.e. tubes) could be extruded under force.

We hypothesized that the combination of support-induced increased membrane viscosity, lipid pinning and membrane stretching all contribute to a membrane state that is permeable to small molecules. Membrane pinning to the substrate^20^ and the associated increase in viscosity^39,72^ both interfere with pore resealing, as recently shown to result in partial membrane coating^94^. Additionally, membrane stretching reduces the energy barrier for pore formation^95,96^, making SLBs more prone to rupture and a state of open pores. It is striking to observe that these effects are also observed in highly packed and mechanically stable Lo membranes. The formed pores are very small and not easily resolvable, the reason why they have remained elusive. Other model systems have been shown to be leaky. Free-standing giant plasma membrane vesicles were shown to be permeable to molecules as large as 40 kDa (∼ 5 nm)^97^. Although the presence of pores may limit the applicability potential of supported membranes as cargo carriers, these pores are very small and larger cargoes could be efficiently retained, with some strategies been employed to coat surface defects, including liposome rupture on uncoated patches^98^.

In the stretched state, SLBs become highly fusogenic, likely due to increased lipid area and enhanced defect formation, achieving complete fusion (i.e., full fusion) in under 7 ms, even for membranes that would not typically fuse. In principle, cushioning SLBs on a polymer support may circumvent some of the limitations of bilayers formed on a solid support, providing a more hydrated and fluid environment. However, these systems too have limitations, such as membrane thickening and partial recovery of lipid diffusion only in a narrow range of polymer density^99^. The findings underscore the utility of both free-standing and supported bilayers as valuable yet fundamentally distinct model systems. We anticipate that the results presented here will have significant impact in the field of nanoparticle-based drug delivery systems as well as in fundamental membrane biophysics.

## AUTHOR CONTRIBUTION

RBL and WHR designed the study. RBL performed and analysed the experiments and wrote the first manuscript draft. WHR edited the manuscript and supervised the study.

### ACKNOWLEDGMENTS

The authors thank Pedro Buzón for his assistance with the optical tweezers, Maisara Elkatan for processing some of the FLIM data and Ceri Richards and Guus van den Borg for their assistance with TIRF experiments. RBL thanks Caio de Jong de Lira for his assistance with the manuscript writing.

## MATERIALS AND METHODS

All materials and chemicals were used as obtained and were not further purified. The phospholipids 1,2-dioleoyl-sn-glycero-3-phosphocholine (DOPC), L-α-phosphatidylserine (Brain, Porcine) (sodium salt) (_brain_PS), 1,2-dioleoyl-sn-glycero-3-phosphoethanolamine (DOPE), and 1,2-dioleoyl-3-trimethylammonium-propane (DOTAP), sphingomyelin (brain, porcine), cholesterol (Chol); the fluorescent dyes 1,2-dipalmitoyl-sn-glycero-3-phosphoethanolamine-N-(lissamine rhodamine B sulfonyl) (ammonium salt) (DOPE-Rh) and 1,2-dioleoyl-sn-glycero-3-phospho-L-serine-N-(7-nitro-2-1,3-benzoxadiazol-4-yl) (ammonium salt) DOPE-NBD, 16:0-06:0 NBD PC (NBD-PC) and 16:0-06:0 NBD PG (NBD-PG) were purchased from Avanti Polar Lipids (Alabaster, AL). Lipid solutions were prepared in chloroform and stored at −20 °C until use. DOPE-Atto647N was purchased from AttoTech h (Siegen, Germany). Flipper-TR plasma membrane (FliptR) was purchased from Spyrochrome (Stein am Rhein, Switzerland). The fluorescent dye KU530 NHS-Ester was purchased from KU dyes (Copenhagen, Denmark). Glucose, sucrose, CaCl_2_, EDTA, and the fluorescent probe Bodipy C_16_ (BODIPY FL C_16_; 4,4-Difluoro-5,7-Dimethyl-4-Bora-3a,4a-Diaza-s-Indacene-3-Hexadecanoic Acid), sulforhodamine B (SRB), Rhodamine 6G, Dextran-FITC 3 kDa, Dextran-FITC 10 kDa and solid (3 μm) and mesoporous (3 μm, ∼ 2 nm pore size) beads were purchased from Sigma-Aldrich (St. Louis, MO, USA). Latex polystyrene beads 3.11 μm were purchased from Spherotech (Lake Forest, IL USA).

GUVs were prepared using the PVA-lipid hydration method^100^. A 2 wt% PVA solution (70-100 μl) in Mili-Q water was homogeneously spread on a glass coverslip and the water was dried by placing the slides on a hot plate at 60-80 ◦C for ∼ 5 minutes to form a semi-dried polymer film. During evaporation, the PVA solution was continuously homogenized on the glass with the help of a pipette tip. Afterwards, a ∼ 10 ml lipid mixture (3 mM concentration) in chloroform was spread on the PVA film and the solvent was evaporated under a argon stream. A coverslip pair was then sandwiched between a Teflon spaced forming a 1.8 ml chamber volume and the slides were closed with clippers. The hybrid film was then hydrated with a 200 mM sucrose solution at room temperature for Ld GUVs, and at 70◦C for Lo GUVs for ∼ 30 minutes, after which they were ready for use. for form the GUVs. For imaging, the GUVs were usually diluted 2-5 fold in isotonic glucose solution for imaging and/or incubation.

Large unilamellar liposomes were prepared using sonication. The equivalent lipids in chloroform were added to the bottom of a glass chamber and chloroform was evaporated under a stream of Argon and further evaporated under vacuum for 1 to 2 h. The lipid film was hydrated with a 200 mM sucrose solution and vigorously vortexed until full film detachment, forming multilamellar vesicles to a final 2 mM lipid concentration. For content mixing experiments, the aqueous fluorescent probes SRB or Dextran 10 kDa were pre-diluted in the hydrating sucrose solution at 50 μM (SRB) or 0.1 mg/ml (Dextran 10 kDa). The multilamellar vesicles were sonicated in a bath sonication for ∼20 min, after which they were ready for use. GUVs and liposomes were incubated by diluting the samples in isotonic glucose. The liposomes were prediluted in sucrose to 100 μM lipid concentration and mixed with 50 μl GUVs in glucose at the desired final liposome concentration for a final 100 μl solution and incubated for 15 min.

Supported lipid bilayers were formed by liposome rupture. Planar SLBs were formed by incubating 500 μM of the respective liposomes (lipid concentration) with plasma-cleaned clean microscope glass slides for 30 minutes. Afterwards, excess non-ruptured liposomes were removed by washing the samples with isotonic liposome-free sucrose solution. For SLBs formed on beads, stock bead solution was vigorously stirred and diluted 100 times, centrifuged (4-6 rcf for 10 minutes). The supernatant was discarded and the beads were resuspended in 200 mM sucrose solution. The resuspended beads were incubated with 500 mM liposomes in the presence of 5 mM CaCl_2_ for 30 minutes. Non-ruptured liposome excess was removed by washing the samples 3 times by centrifugation (4-6 rcf for 10 minutes), in which the discarded supernatant was being replaced by isotonic sucrose. For Ld membranes, the SLBs were prepared at R.T. whereas for SM:Chol (7:3, mole ratio) liposomes in the Lo phase, SLBs were prepared at 60-70 C.

Wide-field fluorescence microscopy was performed on a AxioObserver (Zeiss) fluorescence microscope. DOPE-NBD and DOPE-Atto647 were excited using a 470 nm and a 625 nm excitation filter and imaging was performed with a 100x (1.4 NA, oil immersion) objective.

Spot-variation fluorescence recovery after photobleaching (sv-FRAP) was performed on a Zeiss LSM 710 confocal scanning microscope based on modifications from previous single-spot FRAP^101^. The samples were imaged through a 40X (1.2 N.A.) W Korr M27 water immersion objective. The fluorescent probes NBD-PC and NBD-PG were excited with a 488 nm argon laser and its emission detected in the range of 500–565 nm. The nominally selected bleaching areas (0.5, 1, 1.5 and 2.5 μm^2^) were used to selectively photobleach the probes in either the GUVs or SLBs. In both cases, the samples were homogeneous at the microscale. The GUVs were immobilized in a 0.3 wt% agarose solution prepared in 200 mM glucose according to prior protocols^57^. The obtained data was analyzed using Fiji (NIH, USA). The half time of recovery t_1/2_ was measured from the FRAP recovery curves and defined as the time at which intensity is half of its final intensity F_1/2_ = (F_o_ + F_∞_)/2, where F_o_ and F_∞_ are the fluorescence intensity in the first post-bleach image and after full recovery, respectively. To account for fluorescence recovery during photobleaching, we used an approach previously reported^102^. We fit a line profile across the bleached area in the first frame after photobleaching (Figure S1) to obtain the effective bleaching radius (r_e_), f(x) = 1-Kexp(−2x^2^/r_e_ ^2^), where K is the bleaching depth. The determined re was used for diffusion coefficient (D) calculation according to D = r_e_^2^+r_n_^2^/8t_1/2_, where r_n_ is the user defined nominal radii.

Fluorescence lifetime imaging microscopy was perfomed on an Microtime200 equipped with time-correlated single-photon counting (PicoQuant) mounted on a inverted Olympus IX73 microscope using the Symphotime64 software. The samples were illuminated with a 100x (1.4 N.A.) oil immersion objective (UPLSAPO, Olympus) and emission collected on the same objective. Fluorescence was filtered from the excitation by band or long-pass filters depending on the dye used. The green dyes FliptR, Bodipy C_16_ and KU530 were excited with a 481-nm laser and their emission was collected using a 525/50-nm band-pass filter, except for FliptR, whose emission was collected using a 582/75-nm band-pass filter. DOPE-Atto647N was excited with a 638-nm laser and its emission collected with a band-pass filter 690/70 nm. The images were acquired and analyzed using the SymPhoTime 64 software and the samples were excited with a pulsed 20-MHz repetition rate as reported in^60^. Unless stated otherwise, the samples were imaged with 128 x 128 pixels, 1-ms dwell-time, and 300 mm/pixels with typical acquisition times of ∼ 40 s. GUV membrane signal at the equator was manually selected for individual GUVs and all pixels used for fitting without binning. The fluorescence decays were fitted using an n-exponential tail fit with a single (n = 1) or bi-exponential (n = 2) decay model with I(t) is the intensity at time t and I_0_ is the intensity at t = 0. A_1_ and A_2_ are pre-exponential factors associated with lifetime components t1 and t2, respectively, and B is the fluorescence background.

Optical tweezers experiments were performed on a commercial setup (LUMICKS, Amsterdam) combining a dual optical trap, a three-colour confocal microscope and a four-channel microfluidic cell^103^. The setup was mounted on an automated XY stage to allow sample positioning. Every trapped bead pair was calibrated for their power-spectrum before the measurements so as to obtain the trapping stiffness. Bead-to-bead distance was determined using bright-field imaging combined with a bead tracking algorithm. A motorized stage was used to control the trapping position inside the microfluidic chamber. The beads were brought to contact for a typical period of 30-60s and forces of 10-30 pN, upon which they were retracted and retraction forces were determined. We observed substantial variability in responses (interacting vs. non-interacting pairs) across experiments, which was independent of both interaction force and contact duration. Bilayer-coated beads of different compositions and labelled with fluorescent probes of different colours were loaded on separate channels and identified from confocal imaging upon 488, 532 or 638 nm excitation for DOPE-NBD, DOPE-lissamine rhodamine and DOPE-Atto647, respectively. The two-dimensional images were obtained from confocal imaging on an area of interest and the fluorescence signal was collected on single-photon avalanche photodiodes.

Total internal reflection fluorescence microscopy was carried out on a home-built TIRF microscope assembled on an IX-7 (Olympus) inverted fluorescence microscope as previously reported^82,104^. Excitation and emission were collected through a 60× oil immersion objective (NA 1.45, Olympus) and the fluorescence signal detected on a EM-CCD camera (Hamamatsu). Sulforhodamine B or DOPE-lissamine rhodamine were excited with a 561 nm laser (Coherent), whereas DOPE-Atto647 was excited with a 638 nm laser. To determine bilayer fluidity, regular fluorescence recovery after photobleaching checks were carried out every experimental day to check bilayer fluidity. Planar supported lipid bilayers were formed on clean glass slides upon incubation and rupture of sonicated DOTAP:DOPE (50:50 mole ratio) liposomes for 30 minutes followed by removed of free liposomes with an excess of solution. The liposomes either contained 0.2 mol% DOPE-lissamine rhodamine or were devoid of any fluorescent dye. For single-fusion experiments, fluorescently-labelled DOPC or DOPC:brains (50:50 mole ratio) sonicated liposomes were introduced into the flow cell at a typical concentration of 4 μM lipids. Fusion events were recorded under exposure times that varied from 7-50 ms/frame and a typical total length of 10000 images per video. The obtained files were processed using Fiji (NIH, USA).

